# Quaddles: A multidimensional 3D object set with parametrically-controlled and customizable features

**DOI:** 10.1101/194381

**Authors:** Marcus. R. Watson, Benjamin Voloh, Milad Naghizadeh, Thilo Womelsdorf

## Abstract

Many studies of vision and cognition require novel three-dimensional object sets defined by a parametric feature space. Creating such sets and verifying that they are suitable for a given task, however, can be extremely time-consuming and effortful. Here we present a new set of multidimensional objects, *Quaddles*, designed for a study of feature-based learning and attention, but adaptable for many research purposes. Quaddles have features that are all equally-visible from any angle around the vertical axis, and which all have similar response biases on a feature detection task, thus removing one potential source of bias in object selection. They are available as two-dimensional images, rotating videos, and FBX object files suitable for use with any modern video game engine. We also provide examples and tutorials for modifying Quaddles or creating completely new object sets from scratch, hopefully greatly speeding up the development time of future novel object studies.

**Author’s note:** This work was supported by grant MOP 102482 from the Canadian Institutes of Health Research (TW) and by the Natural Sciences and Engineering Research Council of Canada Brain in Action CREATE-IRTG program (MRW, TW). The funders had no role in study design, data collection and analysis, the decision to publish, or the preparation of this manuscript. Authors would like to thank Hongying Wang for technical support, and Isabel Gauthier for comments on a draft version of the manuscript.

The study described herein was approved by the York University Office of Research Ethics (Certificate # 2016-214).

A draft version of this manuscript is available online at BioR_χ_iv:

## Introduction

In many experiments in the cognitive sciences, participants must view three-dimensional stimuli, or two-dimensional projections of three-dimensional stimuli, that they have not encountered before. Such novel object sets have been used in studies of such phenomena as object recognition and discrimination (e.g. Biederman & Gerhardstein, 1993; Bülthoff & Edelman, 1992; Chuang, Vuong, & Bülthoff, 2012; Gauthier, James, Curby, & Tarr, 2003; Harman, Humphrey, & Goodale, 1999; Harman & Humphrey, 1999; Hayward & Tarr, 1997; Richler, Wilmer, & Gauthier, 2017; Tarr, Bülthoff, Zabinski, & Blanz, 1997; Wong & Hayward, 2005), perception and attention to different object properties (Arnott, Cant, Dutton, & Goodale, 2008; Cant & Goodale, 2007), memory for objects (Humphrey & Khan, 1992; Knutson, Hopkins, & Squire, 2012; Mercer & Duffy, 2015), facial perception and recognition (e.g. Gauthier & Tarr, 1997; Gauthier, Tarr, Anderson, Skudlarski, & Gore, 1999; Wong, Palmeri, & Gauthier, 2009), category representation (e.g. Wallraven, Bülthoff, Waterkamp, van Dam, & Gaissert, 2014; Williams, 1998), conditioned fear responses (e.g. Barry, Griffith,Vervliet, & Hermans, 2016; Bennett, Vervoort, Boddez, Hermans, & Baeyens, 2015; Scheveneels, Boddez, Bennett, & Hermans, 2017), linguistic demonstratives and gestures (Cooperrider, 2015), and emotional influences on perception (Estes, Jones, & Golonka, 2012). Some sets of novel object have also been presented in papers specifically written to encourage their adoption by other researchers (Barry, Griffith, De Rossi, & Hermans, 2014; Buffat et al., 2014), or simply hosted online (Harris, n.d.) for other researchers to use. Figure 1 shows representative exemplars of these sets.

**Figure 1:**
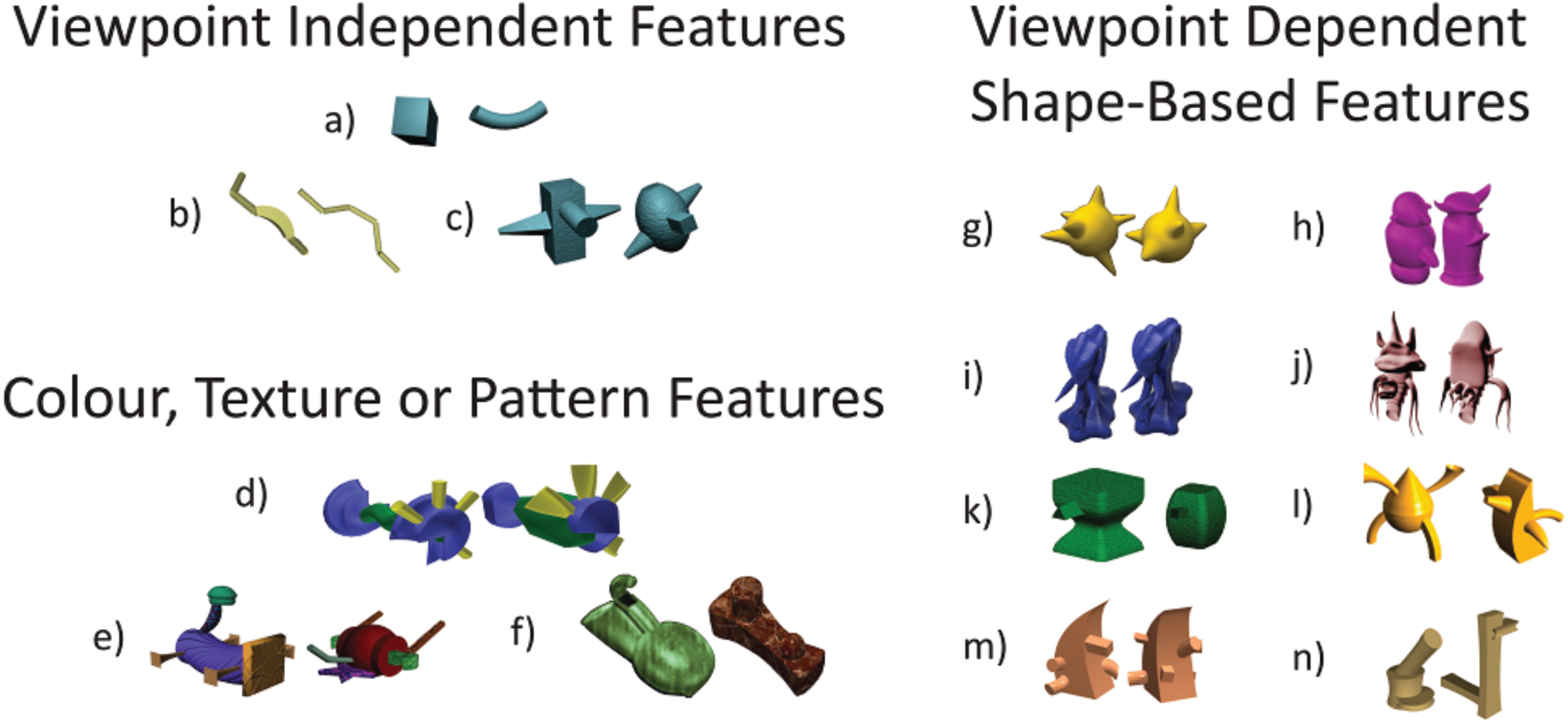
Representative exemplars from a number of previously-reported sets of novel objects. Most do not have features that are viewpoint-independent, or that are defined by elements other than shape, both of which we needed for studies with multidimensional view invariant features. No set has both. a) Geons (Biederman & Gerhardstain, 1993; Hayward & Tarr 1997), b) Strings (Biederman & Gerhardstein, 1993; Bulthoff & Edelman, 1991; Tarr et al, 1997), c) Multi-Geons (Biederman & Gerhardstein, 1993),d) Yadgits (Harris, n.d.), e) Fribbles (Barry et al 2014; Williams 1998); f) Nonsense objects (Cant & Goodale, 2007; Humphrey & Khan, 1992); g) Ameoboids (Bulthoff & Edelman, 1991; Hayward & Wong, 2005), h) Greebles (Gauthier & Tarr, 1997), i) Yufos (Gauthier et al., 2003), j) Sheinbugs (Richler et al, 2017), k) Two-Part Objects (Hayward & Tarr, 1997), l) Pair-wise similar objects (Biederman & Gerhardstein, 1993), m) Geons with occlusion (Hayward & Wong, 2005), n) Ziggerins (Wong et al., 2009). Exemplar pairs g), m), and n) courtesy of Michael J. Tarr, Center for the Neural Basis of Cognition and department of Psychology, Carnegie Mellon University, http://www.tarrlab.org, a), b), c), d), e), h), i), k) & l) were taken from http://wiki.cnbc.cmu.edu/Novel_Objects, j) was taken from http://gauthier.psy.vanderbilt.edu/resources/, and f) was taken from Cant & Goodale, 2007. Some stimuli were edited to remove background colour in figure.

As the use of novel objects in research has become more commonplace, there has been a parallel rise in studies where participants engage and interact with complex, continually-changing virtual environments. Such dynamic tasks, presented on traditional monitors or stereoscopic displays, enable the presentation of much richer stimuli and the collection of much richer data streams than more traditional static tasks. They have been used to investigate the processes underlying phenomena such as the mechanisms of spatial navigation in humans and other animals (Bohil, Alicea, & Biocca, 2011; Ekstrom et al., 2003; Weisberg, Schinazi, Newcombe, Shipley, & Epstein, 2014), multisensory integration in the determination of one’s own location (Ehrsson, 2007; Lenggenhager, Tadi, Metzinger, & Blanke, 2007), memory retrieval (Watrous, Tandon, Conner, Pieters, & Ekstrom, 2013), priority in attention, gaze, and memory (Aivar, Hayhoe, Chizk, & Mruczek, 2005; Jovancevic, Sullivan, & Hayhoe, 2006), the temporal organization of gaze in realistic tasks (Johnson, Sullivan, Hayhoe, & Ballard, 2014), subliminal cueing (Aranyi et al., 2014; Barral et al., 2014), and the development of brain-computer interfaces (Leeb et al., 2007).

Both dynamic tasks and novel 3D object sets, then, have become standard tools in the cognitive science repertoire. We are not aware of any published work that combines the two, but we anticipate that this will rapidly become commonplace, as more researchers become aware of the power and flexibility these tools enable without a corresponding sacrifice in experimental control. Our laboratory has begun running such studies, in which we examine attentional and oculomotor changes as participants learn about a novel object set in a dynamic environment (Watson, Voloh, Naghizadeh, Chen, & Womelsdorf, 2017). With so many sets of novel objects freely available (see Figure 1), it came as a surprise that we could not find a multidimensional set that met our requirements. Instead, we had to design our own and test their suitability for our task, a much more difficult and time-consuming project than originally anticipated, and one which we hope to make substantially easier for future researchers.

In this paper, we review this novel object set, named *Quaddles* in reference to the four feature dimensions that define the object space. In addition, we describe (and provide links to) tools that enable researchers to design their own parametric object sets quickly and relatively easily. Finally, we present the results of a feature detection task verifying that there are no strong response biases towards any feature values of Quaddles across individuals.

### Introducing Quaddles

The experimental task for which Quaddles were designed has participants moving freely using a joystick around a realistic virtual three-dimensional environment, and choosing between objects in that environment. Their object selections are either rewarded or not, based on the particular feature values of the objects, and they have to learn through trial and error which feature values are associated with reward. Our requirements for these objects were that they have:

- An aesthetically-pleasing appearance.
- Multiple feature dimensions, including non-shape dimensions.
- Multiple feature values along each feature dimension.
- Feature values that do not produce strong response biases across individuals.
- Vertical symmetry
- Features that can all be clearly and simultaneously viewed from any angle around the vertical axis.
- The ability to be exported to any commonly-used image format (PNG, GIF, JPG, etc).
- The ability to be exported to any common video game engine (Unity 3D, Unreal Engine, etc)

There is no set of novel objects we are aware of that meets all these criteria. Very few of these sets have viewpoint-independent features that can all be viewed simultaneously from any angle around the object, and most have purely shape-based feature dimensions (see Figure 1). Furthermore, we are only aware of one object set for which feature similarity has been quantified (Barry et al., 2014). Hence the need for a new set of objects.

Quaddles are defined by four feature dimensions: body shape, arm angle, surface colour, and surface pattern, each of which has two possible values (Figure 2). The set can easily be extended by adding new feature values (Figure 3), by using intermediate (morphed) values between existing values (Figure 3), or by adding new dimensions entirely (Figure 4). All Quaddles shown in these figures are freely available on our website (http://accl.psy.vander-bilt.edu/resources/analysis-tools/) in FBX file format. High definition pictures of each object from a variety of perspectives are also available in JPG formats with black, white, or grey backgrounds, as well as in PNG format with transparent backgrounds. Videos of the objects rotating against black, white and grey backgrounds are also available in mp4 format. Finally, the same website contains a detailed manual and scripts to generate all objects and files, which can easily be modified to create custom object sets.

**Figure 2:**
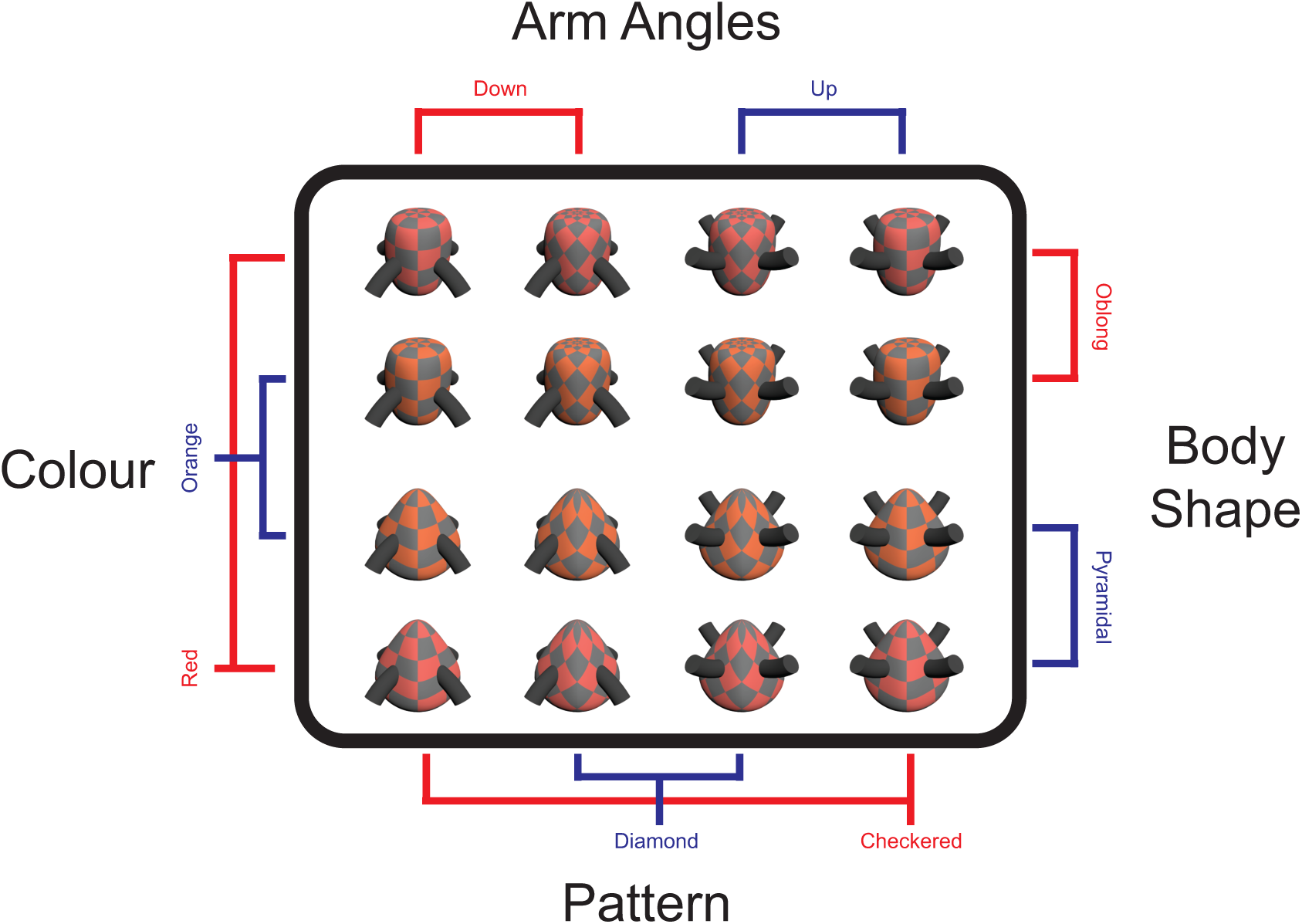
The primary set of 16 Quaddles used in the feature identification study described in this paper.

**Figure 3:**
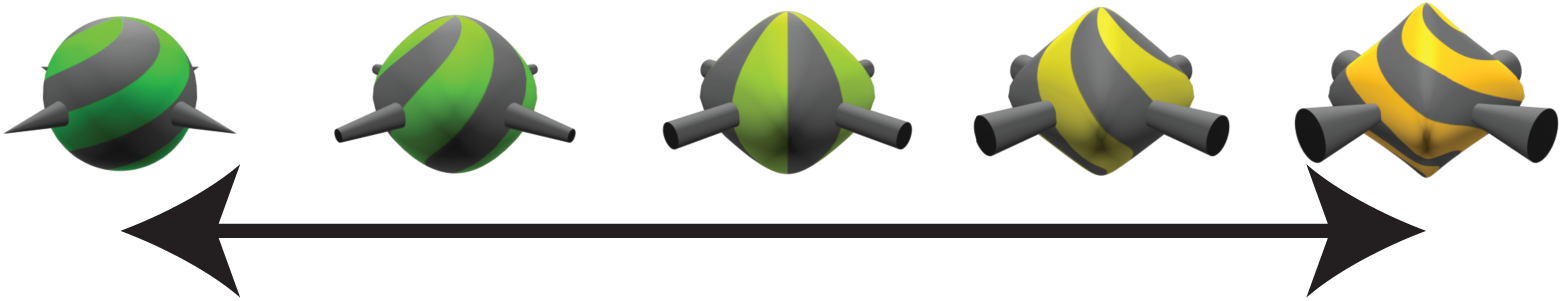
Examples of morphing, using 2 different feature values on each of the four feature dimensions

**Figure 4:**
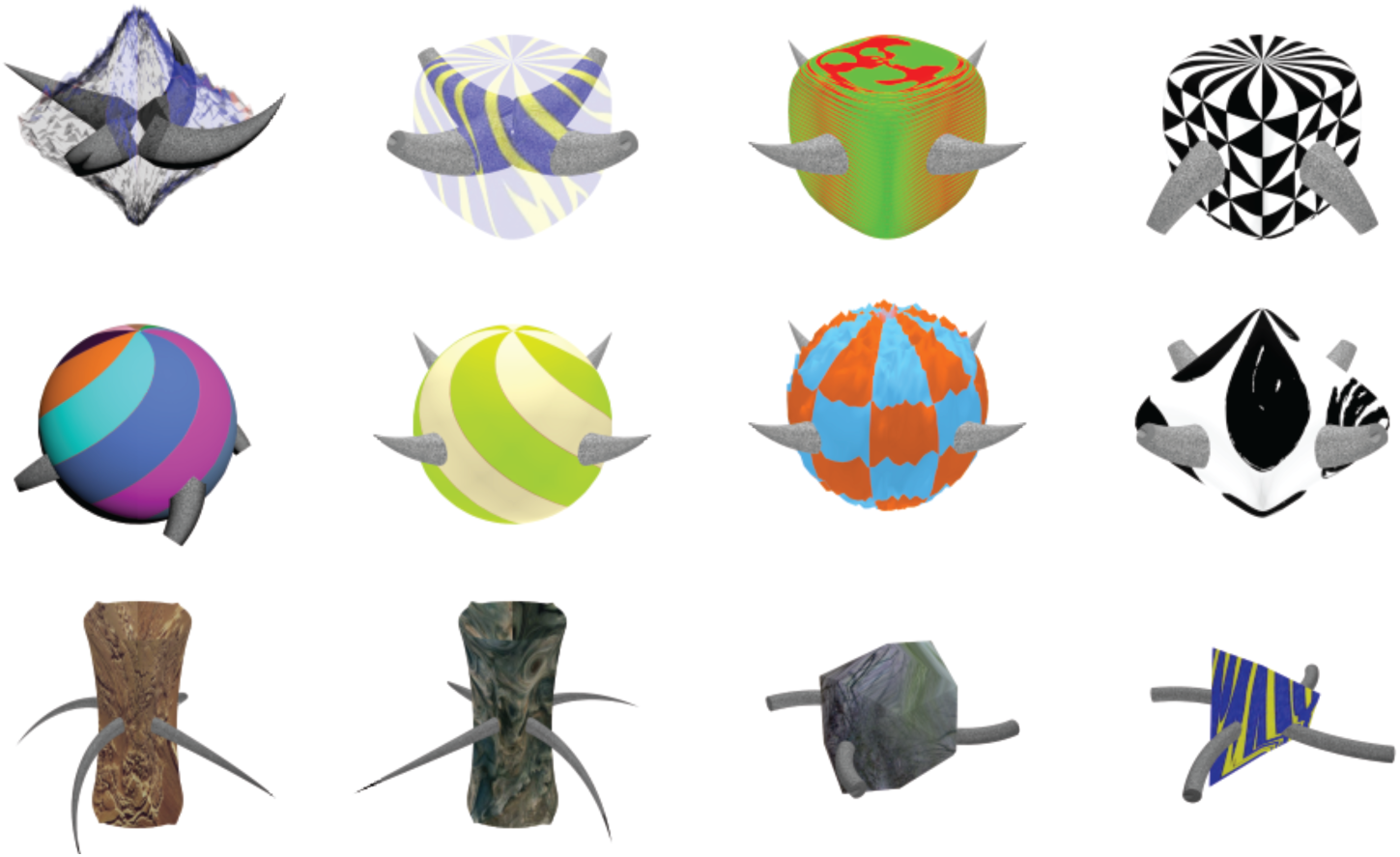
Additional objects showing some possible variations on the basic Quaddle body plan that can be easily generated via scripting.

In the remainder of the paper, we summarize the methods for creating Quaddles (more complete details can be found in the manual, hosted at http://accl.psy.vanderbilt.edu/re-sources/analysis-tools/), and present the results of a feature detection study quantifying response biases to the different feature values. In this task, subjects are cued with two feature values prior to being shown a single Quaddle that contains only one of these values, and have to report which of the two values is present. By pooling response times and accuracies across trials where a given feature value is cued (either validly or invalidly), the response bias to this feature value can be quantified. Furthermore, the stability of these biases can be quantified using consistency measures, both across blocks within individual participants, and between participants. To prefigure our results, this analysis demonstrates that feature value biases are fairly consistent within individual participants, but substantially weaker across participants. Furthermore, by quantifying between-participant biases, these can be used as covariates in analyses of other tasks using Quaddles, thus controlling for their effects.

## Methods

### Stimulus generation

Quaddles were generated using Autodesk 3DS Max software. The primary set, used to generate all results presented in this paper, is defined by four feature dimensions (body shape, branch angularity, pattern, & colour), each of which can take on two possible feature values (e.g. body shape can be pyramidal or oblong), giving a total of 8 feature values and 16 possible objects (Figure 2a).

Textures, which define the surface colors and patterns, are imported from PNG files created in a Matlab script. The neutral gray colour is the same for all objects, while the other colours are chosen within the Cie L^*^c^*^h^*^ space such that L^*^ and c^*^ values (luminance and saturation, respectively) are held constant, but h^*^ values (hue) vary by 15°, meaning that there is a small difference in hue between the two colours, but not in other components of colour. Textures are applied to object surfaces using standard UV mapping options: a spherical wrap for pyramidal bodies and a cylindrical wrap for oblong ones (different wraps were chosen because they resulted in smaller artifacts at the top and bottom of objects).

Quaddle bodies are initially generated as spheres and then moulded into the desired body shapes using free-form deformation (Sederberg & Parry, 1986), in which a lattice of control points is added to the object and manipulated to create the desired shape. Thus, all 4 body shapes are morphs of each other, allowing for intermediary shapes as desired. Each Quaddle has four arms, initially generated as straight cylinders and then morphed into the desired shape, thus also supporting intermediary values. The same is true of both the hues used to define object colour, and their surface patterns. This means it is easy to create objects chosen from anywhere within the feature space defined by the 4 feature dimensions (Figure 3). Given any two objects, one can also create videos of the morph between them, or even objects that morph in realtime in a 3D environment.

Object generation was automated using a 3DS Maxscript which creates and saves complete object sets. An optional function allows JPEG, PNG or other image files to be generated of every object created from any distance, height, and rotation. Experimenters also have the option of saving videos of the objects rotating 360 degrees from any perspective.

For illustration purposes, we generated 2 more feature values along each dimension, and generated partial morphs of the objects along each dimension (Figure 3). We also generated a number of further symmetrical objects using a similar 4-arm and symmetrical body layout, and various different textures and shapes (Figure 4). Making new Quaddles in this way is quite easy using simple modifications of our existing scripts, allowing the powerful and flexible generation of new object sets.

### Experimental procedures

The York University Office of Research Ethics approved the present study as confirming to the standards of the Canadian Tri-Council Research Ethics guidelines (Certificate # 2016-214). Ten participants (mean age 28±3.8 SE; 4 female) ran in the study. One was excluded from further analyses due to chance performance. They were seated approximately 60 cm from a LED monitor with a 60 Hz refresh rate, with heads unrestrained. The entire study, including an instructional tutorial, took approximately 1 hour. The task was coded in the Unity game engine.

For the duration of the experiment, participants viewed a diamond-shaped arena from one of its vertices (Figure 5). At the start of each trial, the floor of the arena changed to one of 20 different textures, chosen at random. These were chosen from a large, free database of textures (https://share.allegorithmic.com), and included a wide variety of hues, contrasts, spatial frequencies, and semantic information. After 200 ms, two cues appeared, each showing an iconic representation of one of the 8 stimulus feature values (two feature values X four feature dimensions). After a further 250 ms (±50 ms jitter), a single Quaddle was displayed at the centre of the screen for 250 ms, subtending 3.5-4.0° visual angle at a 60 cm viewing distance. Subjects had to quickly decide whether the single Quaddle contained the feature value of the left or right iconic image cues. A mask pattern was then flashed over the Quaddle location for 50 ms, after which both it and the Quaddle were removed. The cues remained on screen until participants responded by pressing either the ‘Z’ or ‘/’ keys on a standard keyboard (indicating that the left or right cue, respectively, was accurate), or 2 seconds had elapsed, whichever was quicker. If participants did not respond within 2 seconds, the game was paused and they were asked to respond more quickly on future trials. After response, feedback was presented for 500-600 ms in the form of a coloured border around the chosen cue, with green indicating correct and red indicating incorrect. Each Quaddle had one of the two cued feature values, but not the other, and participants were tasked with reporting which of the two cues was present. After feedback, the cues were removed and the next trial started immediately (see Figure 5).

**Figure 5:**
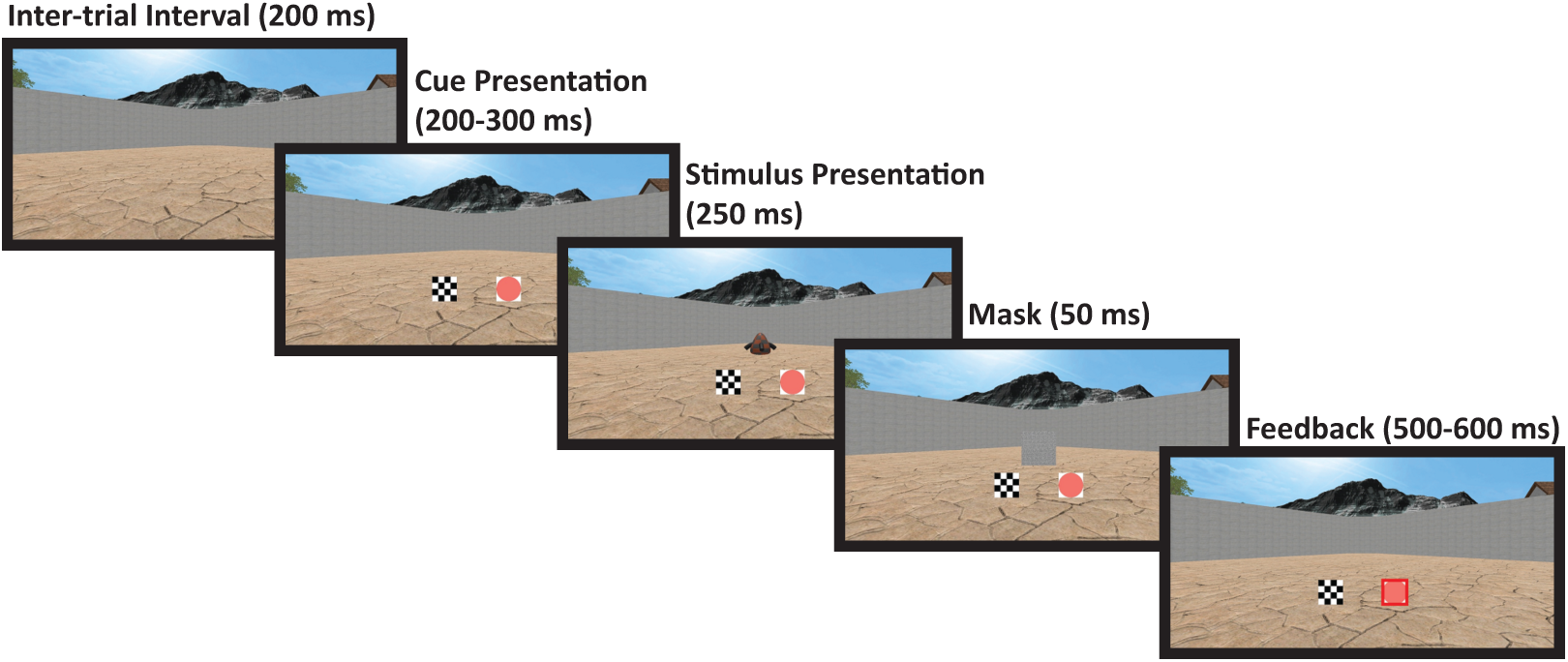
Overview of a feature detection trial. On this trial, the participant incorrectly responded that the presented Quaddle had a reddish colour, instead of a checkered pattern, and so their incorrect choice was outlined in red during the feedback phase. Had they chosen correctly, their choice would have been outlined in green.

Participants were given approximately 5 minutes of training on a slower version of the task prior to starting, and were instructed to respond as quickly and accurately as possible. A single block contained 512 trials, consisting of 32 trials for each of the 16 Quaddles. Each of the 4 feature values present on a given Quaddle was presented as a valid cue 8 times, twice with each of the 4 feature values not found on the same Quaddle as the invalid cue, both on the left and right side. Every 32 trials, each of the 16 Quaddles was shown twice, one with the valid cue on the right, and once on the left, but in all other respects cues, Quaddles, and the side of the valid cue were randomized. After a block, participants were given an optional break. Most participants ran through 3 blocks in approximately 60 minutes, but 3 were only able to finish 2 due to time constraints.

Several pilot versions of the study were run. After each, we adjusted object feature values to try and eliminate any gross response biases. We present the results only for participants run using the final set of feature values, which had the most unbiased performance across feature dimensions.

## Results

One participant was excluded from analyses due to chance accuracy. For the remaining participants, all trials where a given feature value was a valid cue were grouped together, as were all trials where it was an invalid cue. This resulted in 16 groups of trials, across which accuracy and response time on correct trials were averaged (Figure 6). Across all participants, median accuracies for all feature values were between 83-94%, and median correct response times were between 750-830 ms. There were substantial inter-individual differences, with individual mean accuracies for particular feature values ranging from 54-99%, and correct response times ranging from 500-1180 ms. Of note here is that there may be a slight bias against the two surface pattern values (checkers and diamonds): presenting them as cues results in the two lowest median accuracies when they are valid, first- and third-lowest median accuracies when they are invalid, and the two slowest median response times in either case. However the medians (and means) across participants of all feature values are within the 95% confidence intervals of all other values.

**Figure 6:**
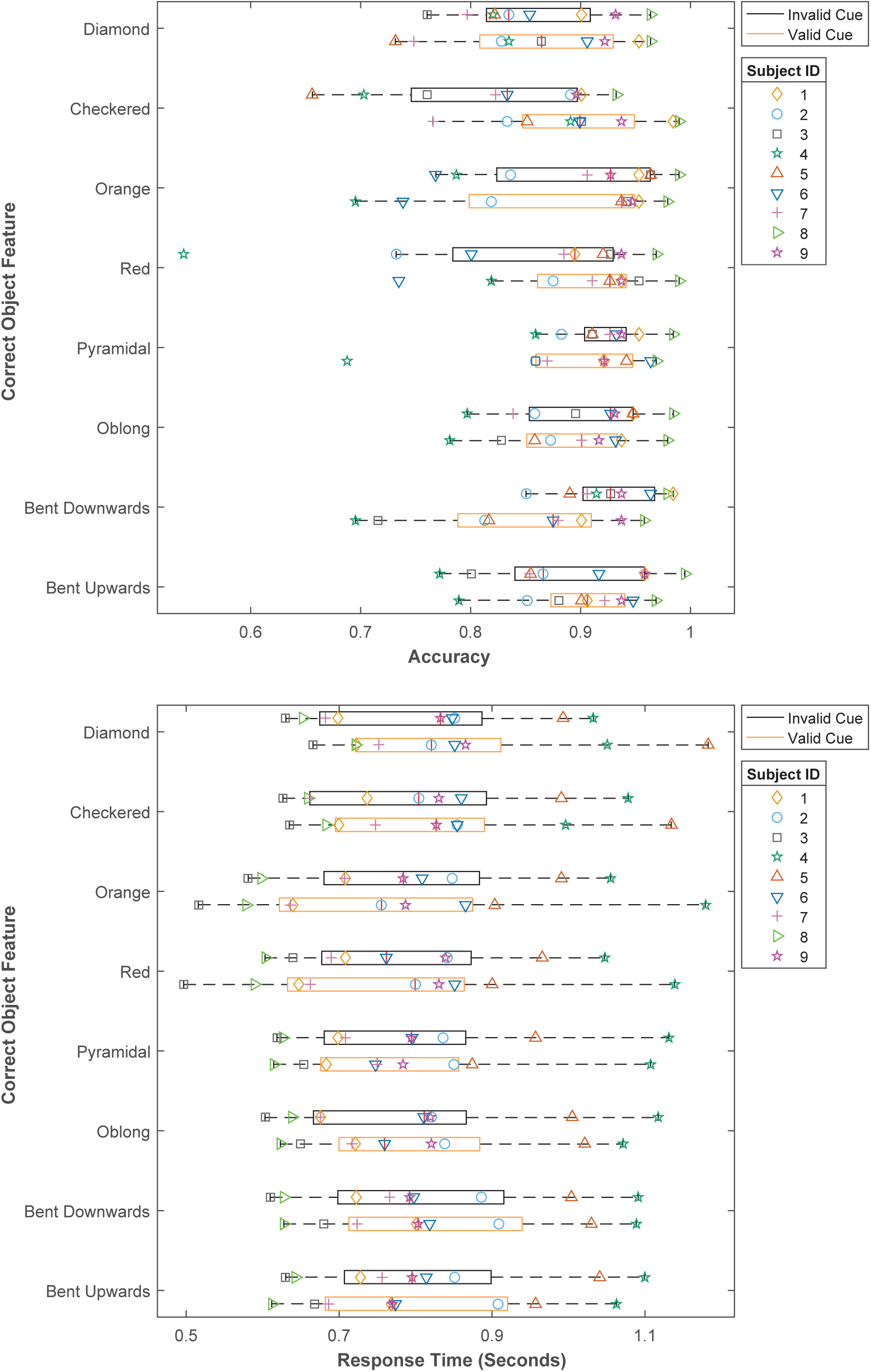
Feature detection study accuracies (b) and correct response times (b) for all trials where a given feature value was presented, either as a valid or an invalid cue.

To more fully characterize response biases, and to control for inter-individual differences in performance, a normalized efficiency score was created from the raw accuracies and re-sponse times. There are 56 cue pairs (8×8 feature values, but the same value cannot be paired with itself), for which accuracies and response times were combined by dividing response time on trials where these cues were presented by the mean accuracy on these trials (Smilek, Enns, Eastwood, & Merikle, 2006; Townsend & Ashby, 1983). Doing this corrects for a speed/accuracy tradeoff in an intuitive way: when accuracy is perfect, efficiency will be identical to mean response time, and as accuracy decreases, the combined score increases, making efficiency scores similar to response time in that smaller values indicate better performance. This assumes that response time and accuracy are linearly correlated, which was supported in the present data (*r* = -0.53, *p* < 0.001). These were then transformed into z-scores using the mean and standard deviation of efficiency scores across all trials for each participant. These 56 scores were combined into 28 for each participant by averaging over the valid-invalid and invalid-valid trials for each feature value pair, also justified on the basis of a strong correlation between the two sets of scores (*r* = 0.83, *p* < 0.001). Finally, these 28 feature value pair scores were combined into 8 feature value scores, by averaging all trials where a given feature value was presented as a cue with any of the other 7 feature values. Each of these scores indicates the bias for a particular feature value, with lower scores indicating greater efficiency. A score of 0 indicates the mean efficiency across all feature values for each participant, and 1 (or -1) indicates a standard deviation away from this mean.

These normalized efficiency scores are shown in Figure 7. Their median values across all participants are close to 0, with no scores more than 0.25 s.d. away from the mean, indicating that most of the variance in response times and accuracies on the present task is due to factors other than response biases to particular feature values. All the 95% confidence intervals for these scores include 0, indicating that there is little, if any, bias for any feature values. However it is perhaps worth noting that the two surface pattern feature values have the two highest scores (0.23 and 0.14), suggestive of a small bias against these values, consistent with the raw data (Figure 6).

**Figure 7:**
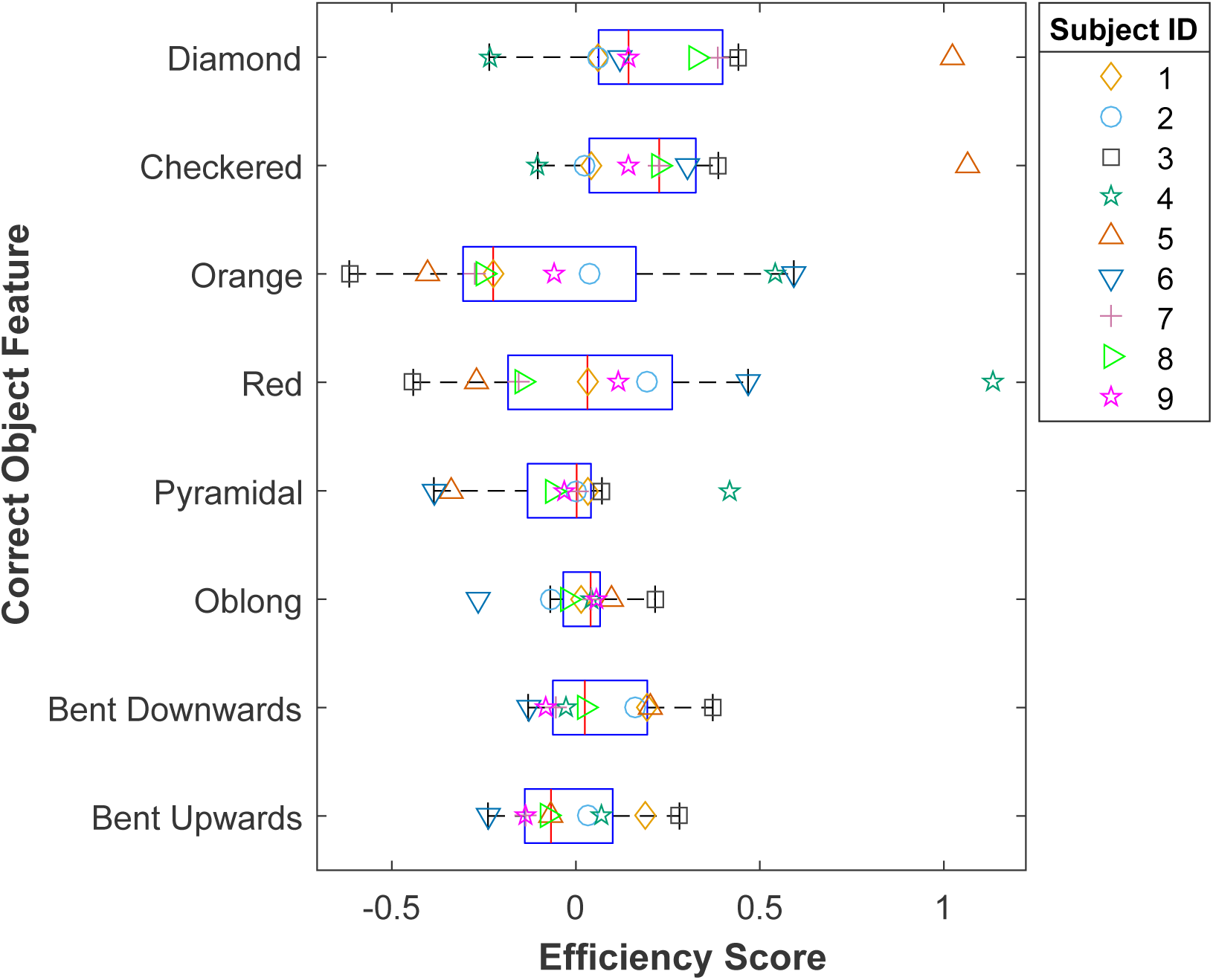
Normalized efficiency scores on the feature identification task. Lower numbers indicate better performance. 0 indicates each participant’s mean efficiency across all trials, and 1 indicates a standard deviation away from this mean.

As well as quantifying the bias for particular feature values, we wanted to quantify how consistent these biases were, both between and within participants. This was accomplished using a tool from the content analysis field, *Krippendorff’s Alpha*, or *Kα* (Hayes & Krippendorff, 2007; Krippendorff, 2011). Kα, which indicates the reliability of multiple sets of scores for a number of items, ranges between -1 to 1, where 1 indicates perfect consistency, 0 indicates a completely random distribution of scores across sets, and -1 indicates perfectly systematic disagreement (Krippendorff, 2008). Generally speaking, Kα is used to measure the consistency of questionnaires or other rating instruments, in which case a high value (e.g.0.80 or higher) is desirable. However in the present case, values approaching 0 indicate a *lack of* consistent bias towards particular feature values, as desired for our object set.

To calculate Kα, normalized efficiency scores were transformed to rank orders. For the between-participants Kα, this was done across all trials to give a single set of scores for each participant. Kα was calculated using a freely-available Matlab script (Eggink, 2012), standard errors and confidence intervals were calculated using a bootstrap method as recommended by Zapf, Castell, Morawietz, and Karch (2016), save that we used 10,000 samples due to the small number of participants, and used bias-corrected and accelerated confidence intervals, which provide more accurate estimates of the true interval (DiCiccio & Efron, 1996). The resulting between-participant consistency is low (Kα = 0.14, SE = 0.17), and its 95% confidence interval includes 0. For the within-participants Kα, we calculated a separate set of efficiency scores for each block performed by each participant, found the rank-ordering of these scores, then calculated a single across-blocks Kα for each participant using these rank orders, and averaged this across participants, using a 10,000 sample bootstrap to calculate standard errors and confidence intervals. This showed that there is a substantial degree of within-participant consistency (Kα = 0.51, SE = 0.12). Thus, individuals have reasonably consistent biases over time, although across individuals these biases are much closer to random (see Figure 8).

**Figure 8:**
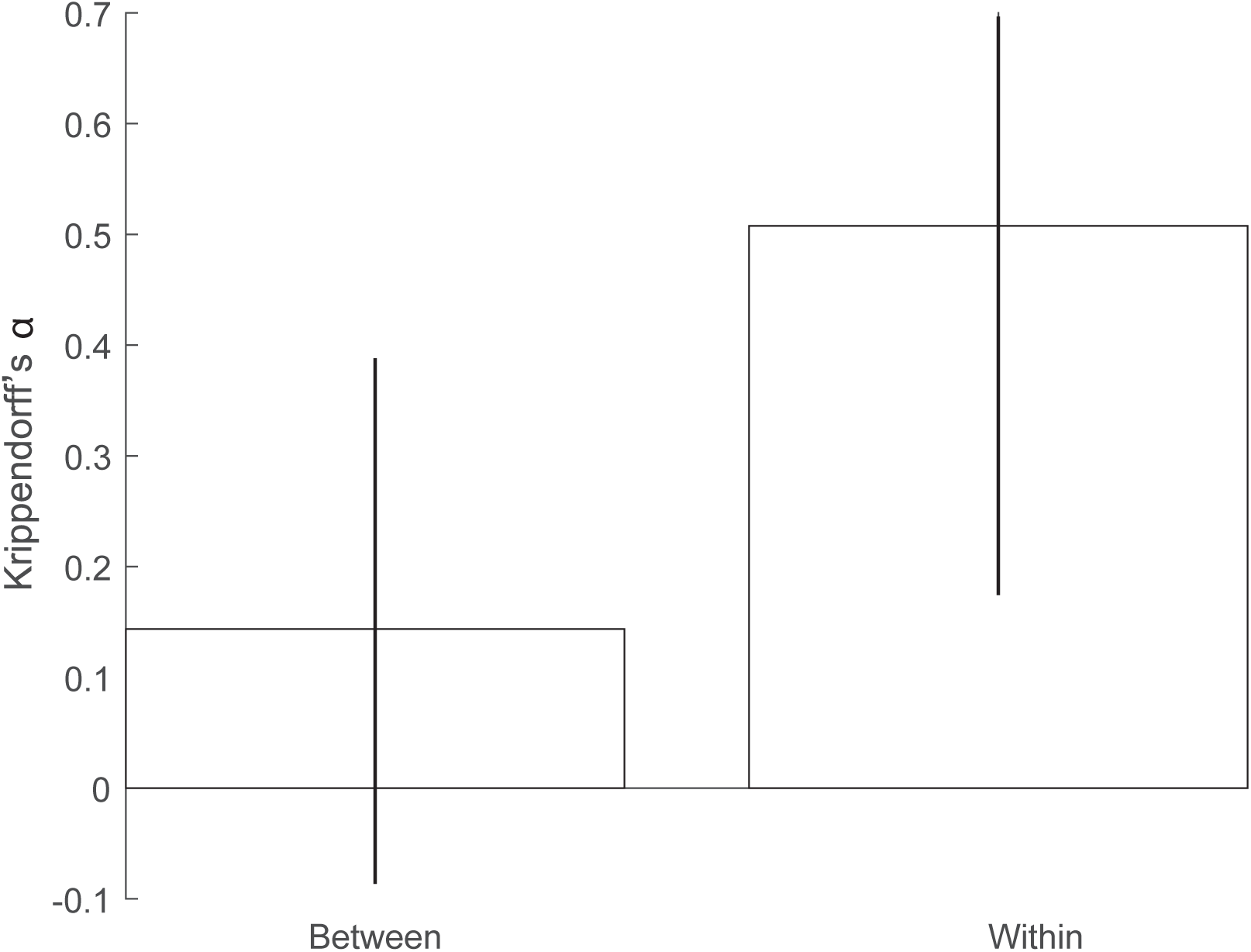
Krippendorff’s alpha values for normalized efficiency score ranks for feature values, between and within participants. Between-partici-pant scores are close to 0, indicating that any given individual’s feature value preferences are close to random. Within-participant scores, on the other hand, are higher, indicating that individual preferences are stable across multiple blocks. Error bars indicate 95% CIs, calculated using 10,000 bootstrap samples.

## Discussion

This paper presented the multi-dimensional set of Quaddle objects, suitable for use both in studies involving navigation through visually-appealing virtual 3D worlds, or for more static studies which require images or videos of multidimensional objects. The results of a simple feature discrimination task show that the feature values along each dimension are roughly equally discriminable (Figure 6), and that while individuals have somewhat stable response biases for particular feature values, these preferences are not strongly shared across participants (Figures 7 and 8). These results suggest that Quaddles can be used “off-the-shelf” in a wide array of tasks that require balanced feature values. They can even be 3D-printed, should an experiment be truly real-world, and can generate stereoscopic images for use with virtual reality or augmented reality experiments.

### Limitations of the discrimination study

While the results of our discrimination task are encouraging, it is important to acknowledge their limitations. First, there is substantial individual variability. For a study where it is critical that each feature value is equally discriminable by all participants (for some arbitrarily small value of “equally”), it might be necessary to produce a much larger set of objects with many intermediate feature values, and run participants on a complex adaptive staircase task (see, e.g., Anderson & Johnson, 2006; Klein & Macmillan, 2001; Kujala & Lukka, 2006; Treutwein, 1999), resulting in a personalized set of objects for each participant. Producing such intermediate objects is possible with relatively simple modifications of our existing scripts. Developing such a staircase task would require careful consideration of the specific requirements of the experiment in question.

Our study presents objects at a single distance. In a study where object distances vary, such as any involving navigation through a 3D world, different features will become more or less discriminable at different distances. Equating discriminability across multiple distances would make for a much longer and more complex discrimination study than was feasible in our time frame.

Finally, our study presented objects within a single arena that does not change, save for the floor, which changes drastically across trials. This is simply because the study for which we developed these objects involves a single arena with floors that change across trials. The surround of an object can have powerful effects on feature discrimination, but our study does not control for these effects, as we reasoned that given the wide variety of floors we present, their effects would be essentially random. Once again, any experimenter for whom this is a critical concern will have to run another set of discrimination studies, modifying the objects and environment as needed.

The general moral here is that *controlling for all factors that affect feature discrimination is not feasible*, because there is no such thing as equal feature discrimination *per se*. We have mentioned three factors that we did not control for, which likely interact in highly non-linear ways: individual preferences, object distance, and visual background. Even if their interactions are completely linear, controlling for all of them simultaneously would require a very complex discrimination task, and a very large number of participants. Furthermore, there are certainly other relevant contingencies that we have not outlined here. Experimenters will have to determine to what degree their particular task requires controlling for different factors that might affect feature value discrimination, and design their objects and discrimination tasks accordingly. Alternatively, instead of controlling for such factors, one could simply quantify their differential effects, and use these as covariates in statistical analyses, to be partialed out from the main effects of the respective studies.

### Possibilities for further customization

Figures 2b, 3 and 4 show several ways in which the basic Quaddle feature space can be manipulated, but there are many other ways in which researchers might change Quaddles for their own purposes. For example, they might wish to remove the vertical symmetry of some, or all, feature dimensions, so that manipulating or navigating around objects would be an important part of identifying them, as is the case with most of the objects shown in Figure 1, as well as with many, if not most, real-world objects. Similarly, it might also be of interest to systematically vary the discriminability or salience of different feature dimensions, and to quantify this variance using a discrimination task. This enables the role of feature bias to be directly studied, as opposed to minimized as with the present object set. Such changes would require trivial modifications to the existing scripts. Aside from these examples, there are of course many other possibilities exist for future studies to implement additional changes.

### Concluding remarks

With this paper, we introduced a new object set, characterized its discriminability and have provided tools to facilitate its use in a wide range of possible future studies. This novel set of 3D objects has normed, parametric features, suitable for a wide range of tasks; the open online access to the examples and tools allows researchers need to rapidly create custom object sets suitable for other studies. This pragmatic aspect resonates well with the spirit of recent toolkits for video game engines that streamline the development and running of dynamic experiments (Doucet, Gulli, & Martinez-Trujillo, 2016; Jangraw, Johri, Gribetz, & Sajda, 2014). The properties Quaddles have make them a suitable set of novel objects for future studies using more realistic and complex tasks, and that the time necessary to develop different objects for such tasks will be significantly reduced.

## References

Aivar, M. P., Hayhoe, M. M., Chizk, C. L., & Mruczek, R. E. B. (2005). Spatial memory and saccadic targeting in a natural task. Journal of Vision, 5(3), 177–193.

Anderson, A. J., & Johnson, C. A. (2006). Comparison of the ASA, MOBS, and ZEST threshold methods. Vision Research, 46(15), 2403–2411. doi:10.1016/j.visres.2006.01.018

Aranyi, G., Kouider, S., Lindsay, A., Prins, H., Ahmed, I., Jacucci, G., … Cavazza, M. (2014). Subliminal Cueing of Selection Behavior in a Virtual Environment. Presence: Teleoperators and Virtual Environments, 23(1), 33–50. doi:10.1162/PRES_a_00167

Arnott, S. R., Cant, J. S., Dutton, G. N., & Goodale, M. (2008). Crinkling and crumpling: An auditory fMRI study of material properties. NeuroImage, 43, 368–378.

Barral, O., Aranyi, G., Kouider, S., Lindsay, A., Prins, H., Ahmed, I., … Cavazza, M. (2014). Covert Persuasive Technologies: Bringing Subliminal Cues to Human-Computer Interaction. In A. Spagnolli, L. Chittaro, & L. Gamberini (Eds.), Persuasive Technology [pp. 1-12]. Springer International Publishing.

Barry, T. J., Griffith, J. W., De Rossi, S., & Hermans, D. (2014). Meet the Fribbles: novel stimuli for use within behavioural research. Frontiers in Psychology, 5, 103. doi:10.3389/fp-syg.2014.00103

Barry, T. J., Griffith, J. W., Vervliet, B., & Hermans, D. (2016). The role of stimulus specificity and attention in the generalization of extinction. Journal of Experimental Psychopathology, 7(1), 143–152.

Bennett, M., Vervoort, E., Boddez, Y., Hermans, D., & Baeyens, F. (2015). Perceptual and conceptual similarities facilitate the generalization of instructed fear. Journal of Behavior Therapy and Experimental Psychiatry, 48, 149–155. doi:10.1016/j.jbtep.2015.03.011

Biederman, I., & Gerhardstein, P. C. (1993). Recognizing depth-rotated objects: Evidence and conditions for three-dimensional viewpoint invariance. Journal of Experimental Psychology: Human Perception and Performance, 19(6), 1162–1182.

Bohil, C. J., Alicea, B., & Biocca, F. A. (2011). Virtual reality in neuroscience research and therapy. Nature Reviews Neuroscience, 12, 752–762. doi:10.1038/nrn3122

Buffat, S., Chastres, V., Bichot, A., Rider, D., Benmussa, F., & Lorenceau, J. (2014). OB3D, a new set of 3D objects available for research: a web-based study. Frontiers in Psychology, 5, 1062. doi:10.3389/fpsyg.2014.01062

Bülthoff, H. H., & Edelman, S. (1992). Psychophysical support for a two-dimensional view interpolation theory of object recognition. Proceedings of the National Academy of Sciences of the United States of America, 89, 60–64.

Cant, J. S., & Goodale, M. (2007). Attention to form or surface properties modulates different regions of human occipitotemporal cortex. Cerebral Cortex, 17, 713–731.

Chuang, L. L., Vuong, Q. C., & Bülthoff, H. H. (2012). Learned Non-Rigid Object Motion is a View-Invariant Cue to Recognizing Novel Objects. Frontiers in Computational Neuroscience, 6, 26. doi:10.3389/fncom.2012.00026

Cooperrider, K. (2015). The Co-Organization of Demonstratives and Pointing Gestures. Discourse Processes, 53(8), 632–656. doi:10.1080/0163853X.2015.1094280

DiCiccio, T. J., & Efron, B. (1996). Bootstrap confidence intervals. Statistical Science, 11(3), 189–228.

Doucet, G., Gulli, R. A., & Martinez-Trujillo, J. C. (2016). Cross-species 3D virtual reality toolbox for visual and cognitive experiments. Journal of Neuroscience Methods, 266, 84–93. doi:10.1016/j.jneumeth.2016.03.009

Eggink, J. (2012). kriAlpha [Matlab script]. Retrieved August 30, 2017 from https://www.mathworks.com/matlabcentral/fileexchange/36016-krippendorff-s-alpha

Ehrsson, H. H. (2007). The experimental induction of out-of-body experiences. Science, 317(5841), 1048. doi:10.1126/science.1142175

Ekstrom, A. D., Kahana, M. J., Caplan, J. B., Fields, T. A., Isham, E. A., Newman, E. L., & Fried, I. (2003). Cellular networks underlying human spatial navigation. Nature, 425(6954), 184–187.

Estes, Z., Jones, L. L., & Golonka, S. (2012). Emotion affects similarity via social projection. Social Cognition, 30(5), 584–609.

Gauthier, I., James, T. W., Curby, K. M., & Tarr, M. J. (2003). The influence of conceptual knowledge on visual discrimination. Cognitive Neuropsychology, 20, 507–523.

Gauthier, I., & Tarr, M. J. (1997). Becoming a “Greeble” expert: Exploring mechanisms for face recognition. Vision Research, 37(2), 1673–1682.

Gauthier, I., Tarr, M. J., Anderson, A. W., Skudlarski, P., & Gore, J. C. (1999). Activation of the middle fusiform 'face area' increases with expertise in recognizing novel objects. Nature Neuroscience, 2(6), 568–573.

Harman, K. L., & Humphrey, G. K. (1999). Encoding 'regular' and 'random' sequences of views of novel three-dimensional objects. Perception, 28, 601–615.

Harman, K. L., Humphrey, G. K., & Goodale, M. (1999). Active manual control of object views facilitates visual recognition. Current Biology, 9, 1315–1318.

Harris, J. (n.d.). Yadgits. Retrieved August 24, 2017 from http://wiki.cnbc.cmu.edu/Novel_Objects

Hayes, A. F., & Krippendorff, K. (2007). Answering the call for a standard reliability measure for coding data. Communication Methods and Measures, 1(1), 77–89.

Hayward, W. G., & Tarr, M. J. (1997). Testing conditions for viewpoint invariance in object recognition. Journal of Experimental Psychology: Human Perception and Performance, 23(5), 1511–1521.

Humphrey, G. K., & Khan, S. C. (1992). Recognizing novel views of three-dimensional objects. Canadian Journal of Psychology, 46(2), 170–190.

Jangraw, D. C., Johri, A., Gribetz, M., & Sajda, P. (2014). NEDE: an open-source scripting suite for developing experiments in 3D virtual environments. Journal of Neuroscience Methods, 235, 245–251. doi:10.1016/j.jneumeth.2014.06.033

Johnson, L., Sullivan, B., Hayhoe, M., & Ballard, D. (2014). Predicting human visuomotor behaviour in a driving task. Philosophical Transactions of the Royal Society of London B: Biological Sciences, 369(1636), 20130044. doi:10.1098/rstb.2013.0044

Jovancevic, J., Sullivan, B., & Hayhoe, M. (2006). Control of attention and gaze in complex environments. Journal of Vision, 6(12), 1431–1450. doi:10.1167/6.12.9

Klein, S. A., & Macmillan, N. A. (2001). Threshold estimation: The state of the art. Perception & Psychophysics, 63(8), 1277–1278.

Knutson, A. R., Hopkins, R. O., & Squire, L. R. (2012). Visual discrimination performance, memory, and medial temporal lobe function. Proceedings of the National Academy of Sciences of the United States of America, 109(32), 13106–13111. doi:10.1073/pnas.1208876109

Krippendorff, K. (2008). Systematic and Random Disagreement and the Reliability of Nominal Data. Communication Methods and Measures, 2(4), 323–338. doi:10.1080/19312450802467134

Krippendorff, K. (2011). Computing Krippendorff's alpha-reliability. Retrieved August 28, 2017 from http://repository.upenn.edu/asc_papers/43

Kujala, J. V., & Lukka, T. J. (2006). Bayesian adaptive estimation: The next dimension. Journal of Mathematical Psychology, 50, 369–389.

Leeb, R., Friedman, D., Müller-Putz, G. R., Scherer, R., Slater, M., & Pfurtscheller, G. (2007). Self-paced (asynchronous) BCI control of a wheelchair in virtual environments: a case study with a tetraplegic. Computational Intelligence and Neuroscience, 79642. doi:10.1155/2007/79642

Lenggenhager, B., Tadi, T., Metzinger, T., & Blanke, O. (2007). Video ergo sum: manipulating bodily self-consciousness. Science, 317(5841), 1096–1099. doi:10.1126/science.1143439

Mercer, T., & Duffy, P. (2015). The loss of residual visual memories over the passage of time. The Quarterly Journal of Experimental Psychology, 68(2), 242–248. doi:10.1080/17470218.2014.975256

Richler, J. J., Wilmer, J. B., & Gauthier, I. (2017). General object recognition is specific: Evidence from novel and familiar objects. Cognition, 166, 42–55.

Scheveneels, S., Boddez, Y., Bennett, M. P., & Hermans, D. (2017). One for all: The effect of extinction stimulus typicality on return of fear. Journal of Behavior Therapy and Experimental Psychiatry, 57, 37–44.

Sederberg, T. W., & Parry, S. R. (1986). Free-form deformation of solid geometric models. SIGGRAPH, 20(4), 151–160.

Smilek, D., Enns, J. T., Eastwood, J. D., & Merikle, P. M. (2006). Relax! Cognitive strategy influences visual search. Visual Cognition, 14, 543–564.

Tarr, M. J., Bülthoff, H. H., Zabinski, M., & Blanz, V. (1997). To what extent do unique parts influence recognition across changes in viewpoint. Psychological Science, 8(4), 282–289.

Townsend, J. T., & Ashby, F. G. (1983). Stochastic modeling of elementary psychological processes. New York: Cambridge University Press. Retrieved from poug

Treutwein, B. (1999). Adaptive psychophysical procedures. Vision Research, 35(17), 2503–2522.

Wallraven, C., Bülthoff, H. H., Waterkamp, S., van Dam, L., & Gaissert, N. (2014). The eyes grasp, the hands see: metric category knowledge transfers between vision and touch. Psychonomic Bulletin and Review, 21(4), 976–985. doi:10.3758/s13423-013-0563-4

Watrous, A. J., Tandon, N., Conner, C. R., Pieters, T., & Ekstrom, A. D. (2013). Frequency-specific network connectivity increases underlie accurate spatiotemporal memory retrieval. Nature Neuroscience, 16(3), 349–356. doi:10.1038/nn.3315

Watson, M. R., Voloh, B., Naghizadeh, M., Chen, S., & Womelsdorf, T. (2017). Information sampling and object selection strategies demonstrate the learning and exploitation of feature relevance. In Neuroscience 2017 Washington, DC: Society for Neuroscience.

Weisberg, S. M., Schinazi, V. R., Newcombe, N. S., Shipley, T. F., & Epstein, R. A. (2014). Variations in cognitive maps: understanding individual differences in navigation. Journal of Experimental Psychology: Learning, Memory, and Cognition, 40(3), 669–682. doi:10.1037/a0035261

Williams, P. (1998). Representational organization of multiple exemplars of object categories. Retrieved August 23, 2017 from http://citeseerx.ist.psu.edu/viewdoc/down-load?doi=10.1.1.5.8336&rep=rep1&type=pdf

Wong, A. C.-N., & Hayward, W. G. (2005). Constraints on view combination: Effects of self-occlusion and differences among familiar and novel views. Journal of Experimental Psychology: Human Perception and Performance, 31(1), 110–121.

Wong, A. C.-N., Palmeri, T. J., & Gauthier, I. (2009). Conditions for face-like expertise with objects: Becoming a Ziggerin expert – but which type? Psychological Science, 20(9), 1108–1117.

Zapf, A., Castell, S., Morawietz, L., & Karch, A. (2016). Measuring inter-rater reliability for nominal data - which coefficients and confidence intervals are appropriate. BMC Medical Research Methodology, 16, 93. doi:10.1186/s12874-016-0200-9

